# CleanRecomb, a quick tool for recombination detection in SNP based cluster analysis

**DOI:** 10.1101/317131

**Authors:** Mark Østerlund, Kristoffer Kiil

**Author notes:** Correspondence Unit of Foodborne Infections, Statens Serum Institut, Artillerivej 5, DK-2300 Copenhagen, Denmark.

## Abstract

We present CleanRecomb, a tool to quickly filter a SNP matrix for likely recombination events.

**Method:** The method evaluates segments with identical SNP profiles over the genome, based on the assumption that SNPs in the absense of recombination events are uniformly distributed across the genome. The method is evaluated on a set of 9 ST200 *E. coli* genome sequences.

**Results:** The detected recombination events coincide with regions of elevated SNP density.

## Introduction

In public health microbiology, the low cost of DNA sequencing has made inference of phylogeny of closely related strains a routine analysis for outbreak detection of bacterial pathogens. Inference of phylogeny using Single Nucleotide Polymorphism (SNP) can be severely distorted by recombination events either between strains within the dataset, or with an unobserved strain. We introduce CleanRecomb, a tool for filtering out spurious SNP events caused by recombination events. CleanRecomb functions under the assumption that mutations giving rise to SNPs are uniformly distributed over the genome, while apparent SNPs caused by recombination events will be located in a bounded region.

Code is available at https://github.com/ssi-dk/CleanRecomb.

## Method

Clean Recomb takes as input a SNP alignment, with the SNPs ordered by genome position (or by contig position in case of a draft reference genome). Seeing the input alignment as an ordered strain x SNP matrix, each column in the matrix is assigned a profile based on the relative conserved/nonconserved status of each strain. These column profiles are made by iterating through the column and comparing the current base to the previous, condensing the column to a repetition list string for each nucleotide. For example AAATTAACA would be encoded to “3:1,2:2,2:1,1:3,1:1”, as would CCCAACCTC. Note that nucleotides are changed to the index of their occurrence in the current profile. Thus the profile express an ordered pattern of relatedness for the sequences in the alignment. Identical profiles results in identical hash values, thus recurring patterns can be counted. We denote a set of adjacent columns with identical profiles as a segment, and a segments likelihood of occurring can be calculated as follows:

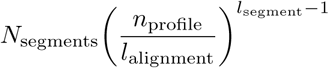

Where *N*_segments_ is the total number of segments, *n*_profile_ is the number of occurences of a given profile, *l*_alignment_ is the number of positions in the SNP matrix, and *l*_segment_ is the length of the current segment with identical SNP profiles.

Segments are evaluated in order of decreasing length, comparing the above statistic to an *α* of 0.05. If a segment fails the threshold, it is marked as a recombination event, and joined into a single point for SNP purposes. Since large recombination events may inflate profile counts and the overall alignment length, both of those are reduced by the size of the discarded segment, and N(segments) is decremented by one, before the next segments are processed. It should be noted that since the input is an alignment of SNPs each position will contain mutations which nullifies any rate of mutation consideration since it would be 100%, thus the simple likelihood model.

## Results

To show an example of the impact CleanRecomb can have we have analysed a set of 9 ST200 *E. coli* sequences using our internal SNP pipeline (https://github.com/PHWGS/ssi-snp-pipeline) and present the maximum parsimony phylogeny before and after applying CleanRecomb.

ST200 shows phylogenetic differences, mainly the 1610T26643 and 1609F54046 branch joins the larger branch. Notably the overall distance of the tree is significantly lowered, indicated by the scale going from 100 to 40 SNPs. As a sanity check for how CleanRecomb marks SNP regions as recombination event we have extracted the SNPmatrix obtained and marked the positions that are filtered.

In Figure 2 the distribution of recombination marked positions can be seen with the relative density of SNP positions on the contig. Density of SNP positions is set as the number of SNPs in ±1000bp from the current, scaled to the highest count observed. It can be seen that recombination and high density mostly coincide. Exceptions include cases such as contig 57 and contig 68, in both cases the entire contig is marked as a single event with low density, this is likely due to plasmid differences or sub-type phylogeny.

**Figure 1:**
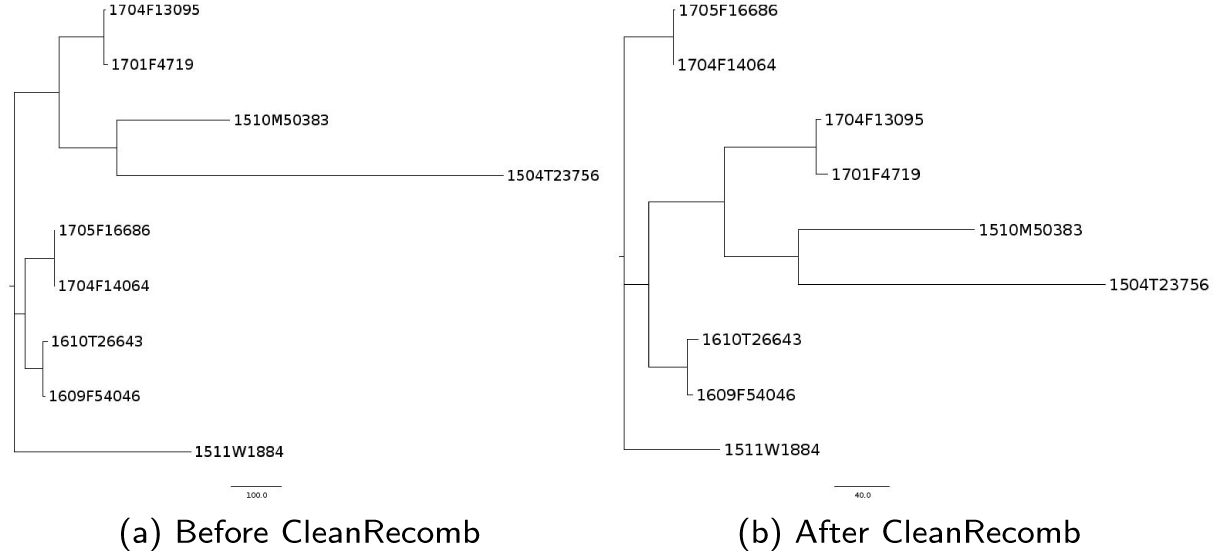
Maximum parsimony tree of 9 ST200 *E. coli* sequences, (a) before and (b) after applying the CleanRecomb method.

**Figure 2:**
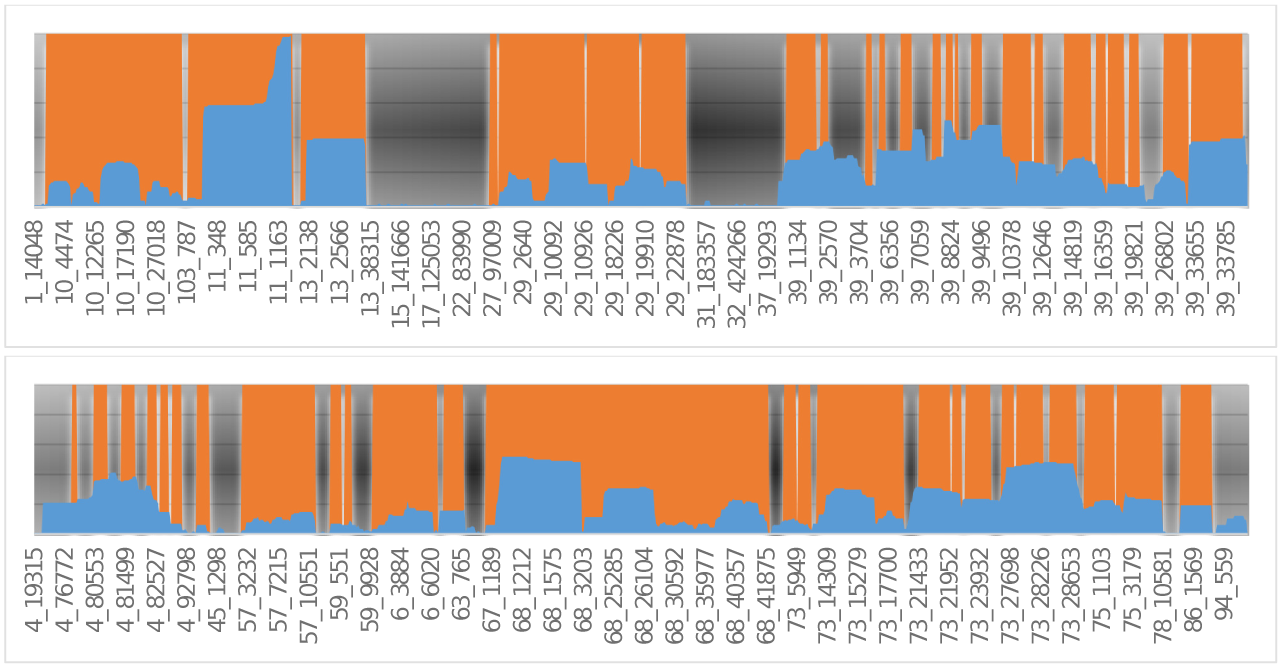
Putative recombination regions and SNP density. Orange regions indicate regions detected by CleanRecomb. Blue bars indicate SNP density in number of SNPs within ±1000bp. Selected SNP positions are marked on the horizontal axis with Contig _position.

## Discussion

If one or more strains have SNPs from random mutation events in the middle of a recombination region the region will be separated into several smaller regions, lowering the power to detect the recombination event. An example of this is contig 29 in Figure 2.

When analyzing closely related strains, the impact is small. If a deep branching strain is included in the analysis, however, this will significantly impact the performance of this method.

Future perspectives: Combining adjacent segments into supersegments might offset the above weakness. Alternatively a sliding window could be used, testing for overrepresented patterns against the binomial distribution.

## Conclusion

Recombination detection is demonstrated to mainly detect regions with a high density of SNPs. While this implementation is aggressively marking SNPs as recombination events it is a useful tool for efficient recombination filtering during SNP analysis as part of pathogen surveillance.

## Author ‘s contributions

Kristoffer Kiil created the algorithm and implemented it in python. Mark Østerlund evaluated the method and drafted the manuscript. All authors proofed the manuscript.

